# Dopaminergic modulation of hemodynamic signal variability and the functional connectome during cognitive performance

**DOI:** 10.1101/130021

**Authors:** Mohsen Alavash, Sung-Joo Lim, Christiane Thiel, Bernhard Sehm, Lorenz Deserno, Jonas Obleser

## Abstract

Dopamine underlies important aspects of cognition, and has been suggested to boost cognitive performance. However, how dopamine modulates the large-scale cortical dynamics during cognitive performance has remained elusive. Using functional MRI during a working memory task in healthy young human listeners (N=22), we investigated the effect of levodopa (L-dopa) on two aspects of cortical dynamics, blood oxygen-level-dependent (BOLD) signal variability and the functional connectome of large-scale cortical networks. We here show that enhanced dopaminergic signaling modulates the two potentially interrelated aspects of large-scale cortical dynamics during cognitive performance, and the degree of these modulations is able to explain inter-individual differences in L-dopa-induced behavioral benefits. Relative to placebo, L-dopa increased BOLD signal variability in task-relevant temporal, inferior frontal, parietal and cingulate regions. On the connectome level, however, L-dopa diminished functional integration across temporal and cingulo-opercular regions. This hypo-integration was expressed as a reduction in network efficiency and modularity in more than two thirds of the participants and to different degrees. Hypo-integration co-occurred with relative hyper-connectivity in paracentral lobule and precuneus, as well as posterior putamen. Both, L-dopa-induced BOLD signal variability modulation and functional connectome modulations proved predictive of an individual’s L-dopa-induced gain in behavioral performance, namely response speed and perceptual sensitivity. Lastly, L-dopa-induced modulations of BOLD signal variability were correlated with L-dopa-induced modulation of nodal connectivity and network efficiency. Our findings underline the role of dopamine in maintaining the dynamic range of, and communication between, cortical systems, and their explanatory power for inter-individual differences in benefits from dopamine during cognitive performance.

## Introduction

Dopaminergic neurotransmission supports cognitive functions, such as flexible updating and stable maintenance of working memory (Goldman-Rakic, 1995; Wang et al., 2004; Vijayraghavan et al., 2007; Cools and D’Esposito, 2011; Kobayashi et al., 2017). Dopamine (DA) plays an important role in modulating synaptic strengths of cortico-striatal pathways that subserve a wide range of cognitive functions (Reynolds and Wickens, 2002). Interestingly, a mere increase of DA level is not beneficial in every individual, and can even be detrimental to task performance depending on individuals’ baseline DA and cognitive performance (Cools and Robbins, 2004; for a review, see Cools and D’Esposito, 2011).

How changing the amounts of DA impacts the cortical dynamics at large scale, and importantly, its relation to individuals’ cognitive performance, has remained unclear. Here, using a double-blind L-dopa vs. placebo intervention during functional MRI, we investigate how DA increase modulates large-scale cortical dynamics during cognitive performance, and whether this modulation can explain intermittent, inter-individually differing behavioral benefits from enhanced levels of DA in healthy young adults.

DA neurotransmission helps maintain the dynamic range of neural circuits by regulating low-frequency tonic firing and phasic activity of DA neurons (Grace, 1995; Venton et al., 2003; Grace, 2016; Kobayashi et al., 2017). This in turn can have important consequences for cortical dynamics at larger scale, and ultimately cognitive performance. Specifically, DA helps maintain moment-to-moment cortical dynamics as measured by signal variability (Garrett et al., 2015; Guitart-Masip et al., 2016), which has been proposed to underlie optimal cognitive performance (McIntosh et al., 2008; Garrett et al., 2015; Guitart-Masip et al., 2016; Armbruster-Genç et al., 2016; see Grady and Garrett, 2014 for a review). Thus, it is plausible to relate the impact of DA on cognitive performance with the modulation of cortical signal variability as a result of changes in neuronal firing patterns (Paladini et al., 2003) or changes in neurovascular coupling and vascular dynamics, which can affect the signal-to-noise ratio of the BOLD signal (e.g., Handwerker et al., 2007; Zaldivar et al., 2014). For instance, compared to younger adults, older adults often exhibited a greater effect of DA challenge on the variability of hemodynamic cortical responses (Garrett et al., 2015; Guitart-Masip et al., 2016), which has been linked to cognitive performance differences between older and younger adults.

Furthermore, the midbrain dopaminergic system innervates widespread areas of cortex ranging from sensory to motor and prefrontal regions (for review see e.g. Jaber et al., 1996; Seger and Miller, 2010; Frank, 2011). Thus, changing DA availability may also modulate brain dynamics on the network level (Kahnt and Tobler, 2017) by altering functional associations among distributed cortical regions, which shape the “functional connectome” in the human brain (Giessing and Thiel, 2012; Carbonell et al., 2014; Finn et al., 2015; Bell and Shine, 2016; Cassidy et al., 2016; Mill et al., 2017). Functional connectivity by definition depends on the statistical associations between brain signals over time. Thus, as higher DA availability may increase moment-to-moment brain signal variability, potentially by enhancing neural phasic activity (Paladini et al., 2003), DA availability could also impact the functional connectivity between widespread cortical regions. Previous accounts based on neural spike measurements suggest that, in the primary visual cortex, much of the variability is shared among large groups of neurons (Lin et al., 2015), and reflects global fluctuations affecting all neurons, which substantially increase correlations among pairs of neurons (Goris et al., 2014; Scholvinck et al., 2015). Thus, a direct relation between higher signal variability and stronger functional connectivity is predictable. Indeed, brain signal variability has been previously suggested as a proxy for information-processing capacity within brain networks (Stam et al., 2002; McIntosh et al., 2008; Lippe et al., 2009; Mišić et al., 2011; Vakorin et al., 2011; McIntosh et al., 2014). Specifically, higher signal variability has been associated with higher nodal centrality (an indication of an important node which has many connections) and network efficiency (an indication of having a higher capacity for parallel information processing; Mišić et al., 2011). Besides, In brain networks derived from neuromagnetic signals, it has been shown that the net information transferred between nodes depends on their signal variability— as measured by sample entropy—and time scale (Vakorin et al., 2011).Accordingly, it has been proposed that brain networks showing higher nodal variability over time have a greater potential for diverse functional configurations (McIntosh et al., 2014). Further, theories of brain metastability suggest that large-scale brain dynamics fluctuate between integrated and segregated network states, where signal variability would facilitate coordinated, flexible shifts between different network configurations (Deco et al., 2011; Tognoli and Kelso, 2014; Deco and Kringelbach, 2017).

Lastly, only a few studies have looked at the impact of DA on the functional connectome. DA (and noradrenaline) appear to increase local functional connectivity within fronto-parietal areas during working memory performance (Hernaus et al., 2017), wheras DA antagonists decrease resting-state network efficiency (Achard and Bullmore, 2007). Nevertheless, it remains unknown how DA modulates large-scale brain network organization during cognitive performance, and how this modulation links to modulations in brain signal variability and behavior.

The current study will address three questions. First, we investigate how changing DA availability modulates large-scale cortical signal and network dynamics during cognitive performance. Second, we ask whether the extent of these modulations can explain the wide range of inter-individual differences in behavioral benefits from DA during cognitive performance (Cools and Robbins, 2004; for a review, see Cools and D’Esposito, 2011). Finally, we examine whether L-dopa-induced modulations in cortical signal variability correlate with L-dopa-induced modulation of the functional connectome. To address these questions, we conducted an fMRI study in which young healthy listeners performed a previously established auditory working memory task (Lim et al., 2015) with and without a single dose (150 mg) of the DA precursor L-dopa. We used graph-theoretical network analysis to explore the impact of L-dopa on the functional connectivity and integration of large-scale cortical networks engaged during the auditory working memory task.

We predicted that L-dopa would increase brain hemodynamic signal variability in cortical regions important for auditory working memory performance. On the network level, our tentative hypothesis was that L-dopa would alter the integration of distributed cortical regions involved in the auditory working memory task, as previous studies suggest higher global integration of brain networks as the system-level mechanism for working memory performance (Cohen and D’Esposito 2016; Finc et al. 2017). Guided by previous work (Cools and Robbins, 2004; Cools and D’Esposito, 2011), we anticipated that the inter-individual differences in behavioral benefits from DA would relate to the degrees of DA modulations in BOLD signal variability and functional networks across participants. Lastly, based on the previous accounts on the role of higher brain signal variability in cortical information processing and flexible network dynamics, we expected a direct relation between DA modulations in BOLD signal variability and the functional connectome.

## Materials and Methods

### Participants

Twenty-two healthy young participants (mean age 27.9 years, age range 25–35 years; 12 females) took part in the study. Two additional participants completed the experiment, but were removed from data analysis due to excessive head movements inside the scanner (i.e., total movement > 3.5 mm of translation or degrees of rotation; scan-to-scan movement > 1.5 mm or degrees). Participants reported no histories of neurological or psychiatric disorders, and none were under any chronic medication. Participants were recruited from the Max Planck Institute for Human Cognitive and Brain Sciences database. Prior to participation all volunteers received a separate debriefing session regarding L-dopa by in-house physicians (B.S. and L.D.). All participants gave written informed consent, and were financially compensated (60€ total). All procedures were in accordance with the Declaration of Helsinki and approved by the local ethics committee of the University of Leipzig (EudraCT number 2015-002761-33).

### Procedure

All participants underwent two double-blind, counterbalanced fMRI sessions, separated by at least one week. Procedures in both sessions were identical. Each session was completed after administering orally either 150-mg L-dopa (Madopar LT; 150-mg Levodopa/37.5-mg benserazide) or placebo. On each scanning (i.e., medication) session, blood pressure and heart rate were measured four times throughout the experiment: before and after in-take of the pills, and before and after the fMRI scanning. Exclusion criteria were based on physiological changes after medication (i.e., greater than ±20 mm Hg in blood pressure and/or ±10 beats per minute in heart rate); none of the participants were excluded based on these criteria. Before and after pill ingestion, participants completed a questionnaire regarding subjective feelings and physical symptoms (Bond and Lader, 1974). None of the participants were excluded due to side effects of drugs. One participant felt nauseated after the completion of the whole experiment (i.e., including the second fMRI experiment); thus N = 22 data were included in the subsequent analyses.

To ensure that fMRI scanning takes place when L-dopa reaches peak plasma concentration (~30 minutes; Dingemanse et al., 1995), the fMRI scan started approximately 35 minutes after in-take of medication. During this interim period (i.e., prior to scanning), participants completed a short practice session in a separate behavioral testing room to ensure that they understood the main experimental (auditory working memory) task. After the practice session, participants were placed in the scanner and went through a short hearing test of the auditory syllables used in the task with the on-going scanner noise in the background. Time from medication administration to scan onset in minutes was later used as a nuisance regressor in all models reported (see *Statistical analysis*).

We acquired eight functional blocks on each medication session (approximately 50 minutes). During each block, participants completed a total of 16 trials. Behavioral responses were collected via MR-compatible response keys. Participants used both index fingers, each assigned to one of two response keys, to give their response. The mapping between hands and response keys were counterbalanced across participants. All auditory stimulation was presented through MR-Confon headphones (Magdeburg, Germany), with Music safe pro earplugs (Alpine Hearing Protection) providing additional attenuation.

### Auditory working memory task

During fMRI acquisition, participants performed a previously established auditory working memory task—a syllable pitch-discrimination task implemented with retroactive cues (see Lim et al., 2015 for full details on the task and materials). Syllable tokens were recorded by a native German female speaker. There were 12 different tokens (with varying pitch) for each syllable category, and one token for each category was randomly selected and presented during the encoding phase. The pitch of syllable tokens was manipulated using Praat (version 5.3). All sound tokens were digitized at 44.1 kHz, had a duration of 200-ms, and normalized to equivalent amplitude (root-mean-squared dB full scale; RMS dBFS).

In brief, on each trial participants encoded two distinct auditory syllables (i.e., /da/ and /ge/ presented in a random order), and detected a change in the pitch of one of the syllables after a delay period (jittered between 9–13 s). During the maintenance of the encoded syllables, one of the two types of retro-cue was presented for 1 s on the screen; on equal probability for each task block, a valid cue (written syllable, “da” or “ge”) or a neutral cue (“xx”) was presented. The valid cue was used to direct participants’ attention to one of the to-be-probed auditory syllable. The neutral cue, however, did not provide any information about upcoming auditory probe. After 5–7 s following the visual retro-cue, an auditory probe syllable was presented.

Within a 4-s time window upon hearing the probe, participants compared the pitch of the probe syllable to that of the same category syllable heard during encoding, and responded “high” or “low” accordingly. Syllable pitch change that occurred at probe was parametrically varied in four steps (±0.125 and ±0.75 semitones) relative to its original pitch heard during encoding. On each trial, participants received a visual feedback (written as “correct” or “incorrect” in German) for 500 ms.

Note that the current study is focused on investigating the overall effect of L-dopa on on-going BOLD signal dynamics during the auditory working memory performance. Thus, we here will not elaborate on the (potentially more complex) phasic effects and interactions involving the cue manipulation.

### MRI data acquisition and preprocessing

Whole-brain functional MRI data were collected using a Siemens MAGNETOM Prisma 3T. Functional data were acquired with a 20-channel head/neck coil using an echo-planar image (EPI) sequence [repetition time (TR) = 2000 ms; echo time (TE) = 26 ms; flip angle (FA) = 90°; acquisition matrix = 64 × 64; field of view (FOV) = 192 mm × 192 mm; voxel size = 3 × 3 × 3 mm; inter-slice gap = 0.3 mm]. Each image volume had forty oblique ascending axial slices parallel to the anterior commissure–posterior commissure (AC–PC) line.

Structural images of fifteen participants were available from the database of the Max Planck Institute (Leipzig, Germany), where a magnetization prepared rapid gradient echo (MP-RAGE) sequence had been used to acquire the structural images [TR = 2300 ms; TE = 2.01–2.98 ms; FA = 9°; 1-mm isotropic voxel; 176 sagittal slices]. For participants without pre-existing structural images in the database, a high-resolution T1-weighted structural image was acquired using an MP-RAGE sequence [TR = 2300 ms; TE = 2.98 ms; FA = 9°; 1-mm isotropic voxel; 176 sagittal slices] at the end of the second fMRI session.

#### Preprocessing

During each functional block 181 volumes were acquired. To allow signal equilibration the first two volumes of each block were removed, and the remaining 179 volumes per block were used for the subsequent analyses. Preprocessing steps were undertaken in SPM12 (Frackowiak et al., 2004). First, the functional images were spatially realigned to correct for head motion using a least square approach and the six rigid body affine transformations. Then, the functional volumes were corrected for slice timing to adjust differences in image acquisition times between slices. This is accomplished by a shift of the phase of the sinusoids that make up the signals. Subsequently, functional images were coregistered to each individual’s structural image. This is achieved by first segmentation of the structural image using tissue probability maps, and then registering the image segments using the rigid body transformations and based on normalized mutual information as an objective function (unified segmentation; Ashburner and Friston, 2005). The resulting functional volumes were then spatially normalized to the standard stereotactic MNI space. No spatial smoothing was applied on the volumes to avoid potential artificial distant-dependent correlations between voxels’ BOLD signals (Fornito et al., 2013; Stanley et al., 2013), which subsequently constitute the input data for functional connectivity analysis (Figure 1C).

**Figure 1.**
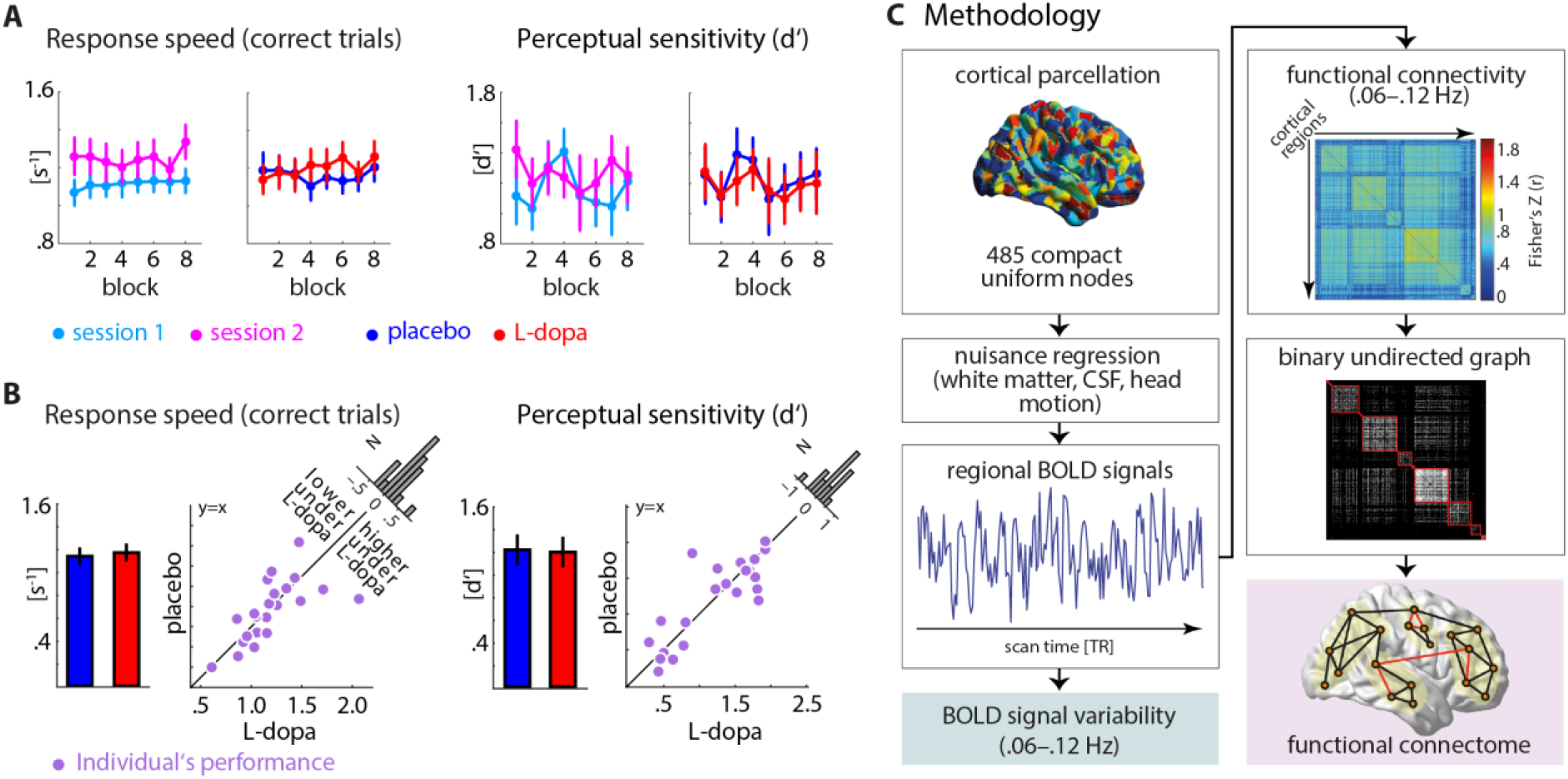
Overview of behavioral performance and methodology. **(A)** Behavioral performances measured by response speed (left) and perceptual sensitivity d’ (right) across eight blocks of the auditory working memory task (adapted from Lim et al., 2015). Block-wise performances are shown based on the order of the scanning sessions and based on the medication sessions. Error bar: ±1 standard error of the mean (SEM). **(B)** Group average and individuals’ behavioral performances in response speed (left) and perceptual sensitivity d’ (right). Scatter plots illustrate individuals’ task performances in the L-dopa and placebo sessions. Histograms (N=22) show the distribution of the modulations (L-dopa minus placebo) of the corresponding measures across all participants. **(C)** Two analysis streams were used to investigate two aspects of brain dynamics: signal variability and large-scale network topology. Both analyses used mean BOLD signals across voxels within each of 485 parcellated cortical regions. BOLD signal variability was computed as the variance of BOLD signal through each fMRI session. The topology of the functional connectome was analyzed using graph-theoretical metrics, which captured network interactions across cortex on local, intermediate and global scales of topology (Betzel and Bassett, 2016). These include local network efficiency (a metric related to the clustering of a network; red triangle), network modularity (a measure of decomposability of a network into communities; light ovals) and global network efficiency (a metric inversely related to the shortest path between nodes; red path).

#### Brain parcellation

Cortical nodes were defined using a parcellated AAL template (Tzourio-Mazoyer et al., 2002) encompassing 485 uniform and compact grey-matter cortical regions (Fornito et al., 2010; Zalesky et al., 2010; Figure 1C). This template was used to estimate the mean BOLD signals across voxels within each cortical region per participant. To examine the robustness of the findings with respect to the definition of network nodes, we also used a functional parcellation encompassing 333 cortical areas (Gordon et al., 2016). In addition, to investigate the significance of striatal contribution in our data, we used a separate functional parcellation of the striatum with seven seed regions (Choi et al., 2012).

To minimize the effects of spurious temporal correlations induced by physiological and movement artifacts, a general linear model (GLM) was constructed to regress out white matter and cerebrospinal fluid (CSF) mean time series together with the six rigid-body movement parameters (Hallquist et al., 2013; Jo et al., 2013). Subsequently, the residual time series obtained from this procedure were concatenated across the eight blocks per medication session for each participant. The time series were further processed through two analysis streams (Figure 1C): one analysis focused on BOLD signal variability (univariate approach) and the other on functional connectivity and network topology (multivariate approach; McIntosh and Misic, 2013; Misic and Sporns, 2016). Since one of the questions that the present study aims to answer is the direct relation between dopaminergic modulation of brain signal variability and networks, we used the same cortical nodes across the two analysis streams.

### BOLD signal variability

Prior to quantifying signal variability, the time series data were band-pass filtered within the range of 0.06–0.12 Hz (see *Functional connectivity* section below for further details). Furthermore, we equated the mean BOLD signal throughout the experiment blocks for each medication session to remove block-wise drifts (e.g., Garrett et al., 2010, 2011, 2013). To this end, we demeaned the residual BOLD time series such that the signal for each cortical region of each block was at zero. For each region, BOLD signal variability was expressed as the variance of the resulting normalized signal concatenated for each medication session (i.e., across 8 blocks). The main focus of the current study is on the amount of behavioral and neural modulations from L-dopa relative to placebo. Instead of computing a difference between two standard deviations of BOLD signal (cf. Garrett et al., 2010, 2011, 2013), the difference measure of the variances was used to compute actual differences in signal variability. Since, given the same amount of difference in two variances, the difference in the corresponding standard deviations tend to be smaller as the absolute variance gets higher, we used variance measures to avoid scaling changes due to the non-linear transformation from variance to standard deviation. The statistical tests were performed using non-parametric procedures (see *Statistical analysis* section), which are robust against non-Gaussian distribution of the signal variability measure.

Signal variability for each cortical region represents region-level signal dynamics. Signal variability on the whole-brain level was computed as the mean variance across all cortical regions (i.e., 485 regions). Signal variability was separately quantified for the L-dopa and placebo sessions. L-dopa-induced modulation of signal variability was then quantified as the difference of BOLD signal variability between L-dopa versus placebo (i.e., L-dopa–placebo). As such, we treated the placebo session as baseline for both brain and behavior during the auditory working memory task.

### Functional connectivity

First, mean residual time series were band-pass filtered by means of maximum overlap discrete wavelet transform (Daubechies wavelet of length 8; Percival and Walden 2000), and the results of the filtering within the range of 0.06–0.12 Hz (wavelet scale 2) were used for further analyses. It has been previously documented that the behavioral correlates of the functional connectome are best observed by analyzing low-frequency large-scale brain networks (Salvador et al., 2005; Achard et al., 2006; Achard et al., 2008; Giessing et al., 2013; Alavash et al., 2015a). The use of wavelet scale 2 was motivated by previous work showing that performance during cognitive tasks predominately correlated with changes in functional connectivity in the same frequency range (Bassett et al., 2010; Alavash et al., 2015b; Alavash et al., 2016). To obtain a measure of association between each pair of cortical regions, Pearson correlations between wavelet coefficients were computed, which resulted in one 485 × 485 correlation matrix (333 × 333 in the case of functional parcellation) for each participant and medication session (Figure 1C).

### Connectome analysis

Brain graphs were constructed from the functional connectivity matrices by including the top 10% of the connections in the graph according to the rank of their correlation strengths (Ginestet et al., 2011; Fornito et al., 2013). This resulted in sparse binary undirected brain graphs at a fixed network density of 10%, and assured that the brain graphs were matched in terms of density across participants and medication sessions (van Wijk et al., 2010; van den Heuvel et al., 2017). To investigate whether choosing a different (range of) graph density threshold(s) affects functional connectivity and network topology differences between placebo and L-dopa (Garrison et al., 2015), we examined the effect of graph thresholding using cost-integration approach (Ginestet et al., 2014; Figure S2).

Mean functional connectivity was calculated as the average of the upper-diagonal elements of the sparse connectivity matrix for each participant per medication session. In addition, three key topological metrics were estimated: mean local efficiency, network modularity, and global network efficiency. These graph-theoretical metrics were used to quantify functional integration of large-scale brain networks on the local, intermediate, and global scales of topology, respectively (Figure 1C; Alavash et al., 2017; Betzel and Bassett, 2016).

For each topological property, we computed a whole-brain metric, which collapses the measure of network integration into one single value, and a regional metric characterizing the same network property but for each cortical region (Rubinov and Sporns, 2010). The regional network metrics were therefore used to localize cortical regions contributing to the L-dopa-induced modulations observed on the whole-brain level. We used nodal connectivity (also known as ‘nodal strength’) as the regional measure of mean functional connectivity. Local efficiency was computed within each cortical region’s neighborhood graph as the regional measure of mean local efficiency. Finally, nodal efficiency was measured to capture the integration of a given region to the entire network, hence representing the regional measure of global network efficiency (see Figure 1C for an illustration). Below we provide the formalization of the above-mentioned graph-theoretical network metrics.

#### Nodal connectivity and degree

Nodal connectivity was measured as the average weights of the connections linking a cortical node to the other nodes. Nodal degree (a simple measure of degree centrality) was quantified as the number of connections per node. Disintegrated nodes were identified as nodes with a degree of zero, accordingly.

#### Global network efficiency

In a graph-theoretical sense, global network efficiency is a measure of information processing capacity of a network. For a given graph *G* comprised of *N* nodes, global efficiency *E*_*global*_ summarizes the capacity of the network for parallel processing across distributed nodes. This metric is estimated by the inverse of the harmonic mean of the shortest path lengths (i.e. the smallest number of intervening connections) between each pair of nodes *L_i,j_*:

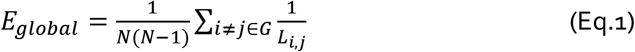

An efficient network is characterized by having a short average minimum path length between all pairs of nodes. Such a network is considered to have high efficiency in parallel (or global) information processing (Bullmore and Sporns, 2009). Likewise, nodal efficiency at node *i*, *E*_*nodal*(*i*)_, (as the regional measure of global network efficiency) is inversely related to the path length of connections between a specific node and the rest of the nodes (Latora and Marchiori, 2001):

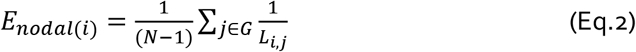

#### Local network efficiency

By slightly zooming out from a given node within a graph, the nearest neighbors of the node that are directly connected to each other form a cluster. This local integration can be quantified based on the local efficiency of node *i*, *E*_*local*(*i*)_ which is mathematically equivalent to global efficiency (Eq.1) but is computed on the immediate neighborhood of node *i*. On the whole-brain level, mean local efficiency can be quantified by averaging local efficiency across all nodes:

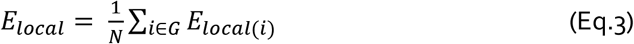

#### Network modularity

Modularity describes the decomposability of a network into non-overlapping sub-networks, characterized by having relatively dense intra-connections and relatively sparse inter-connections. Rather than an exact computation, modularity of a given network is estimated using optimization algorithms (Lancichinetti and Fortunato, 2009; Steinhaeuser and Chawla, 2010). The extent to which a network partition exhibits a modular organization is measured by a quality function, the so-called *modularity index* (*Q*). We used a common modularity index originally proposed by Newman (2006), and employed its implementation in the Brain Connectivity Toolbox (Rubinov and Sporns, 2010) which is based on the modularity maximization algorithm known as Louvain (Blondel et al., 2008). The modularity index is defined as:

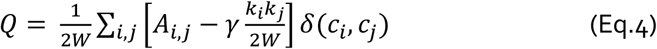

*Q* ranges between −1 and 1. In Eq. 4, *A_i,j_* represents the weight (zero or one if binary) of the links between node *i* and *j*, 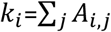 is the sum of the weights of the links connected to node *i*, and *c_i_* is the community or module to which node *i* belongs. The *δ*-function *δ*(*u*,*v*) is 1 if *u* = *v* and 0 otherwise, and 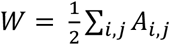. Similar to previous work (Bassett et al., 2010; Alavash et al., 2016), the structural resolution parameter *γ* (see Fortunato and Barthelemy, 2007; Lohse et al., 2014) was set to unity for simplicity. The maximization of the modularity index *Q* gives a partition of the network into modules such that the total connection weight within modules is as large as possible, relative to a commonly used null model whose total within-module connection weights follows 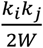. Thus, a “good” partition with *Q* closer to unity gives network modules with many connections within and only few connections between them; in contrast, a “bad” partition with *Q* closer to zero gives network modules with no more intra-module connections than expected at random (Good et al, 2010). Thus, higher *Q* reflects higher functional segregation on the intermediate level of network topology (Rubinov and Sporns, 2010; Betzel and Bassett, 2016). Due to stochastic initialization of the greedy optimization, the module detection algorithm was applied 100 times for each brain graph, and the partition that delivered the highest *Q* value was kept (see Lancichinetti and Fortunato, 2012; Bassett et al., 2013, for other approaches to construct representative high-modularity partitions).

### Statistical analysis

#### Control of medication-session order and other confounding factors

In the present study, all participants went through the same procedure twice on separate days. We did not expect to observe learning during the auditory working memory task, but we examined the potential learning effect within and between the two sessions in order to control for potential confounds in interpreting the L-dopa effect. First, we examined a potential learning over task blocks within each scan session, and whether the degree of learning differs between the two sessions (see Figure S6). Neither behavioral measures showed any significant learning effect across blocks (Table S1). Also, we did not find any significant differences in the degree of potential learning between the first versus second session or between L-dopa versus placebo (Figure 1A; Figure S6; Table S1). Using a similar approach, we investigated changes in measures of brain dynamics within each session over task blocks. We only found significant linear increases in signal variability and functional connectivity over task blocks (see Figure S6 and Table S1 for details). However, important for the current study, there was no significant difference between placebo and L-dopa sessions.

Next, we examined if there is any difference in mean performances between two sessions. We observed significant effects of session order (first vs. second scan session), regardless of medication conditions, on both behavioral and brain measures. Participants’ responses were significantly faster in the second session (F_1,21_ = 8.66; p = 0.008). Also, global network efficiency and network modularity (*Q*) were significantly higher in their second session (*E*_*global*_: Cohen’s d = 0.4, p = 0.008; *Q*: Cohen’s d = 0.43, p = 0.02). This could be arguably due to differences in arousal and/or having an easier time in performing the auditory working memory task, as participants are likely to be more relaxed and adapted to drug administration, scan procedure, and task during the second session (Fan et al., 2012; Kitzbichler et al., 2011; Stevens et al., 2012).

Importantly, to control for potential confounding factors in the interpretation of L-dopa effects, all ensuing comparisons of L-dopa versus placebo were performed on residualized measures, that is, after regressing out covariates of no interest. These covariates included medication-session order (L-dopa first vs. placebo first), body mass index (BMI), and time from medication administration to scan onset.

In recent years, the potential confounding impact of head motion or non-neural physiological trends on the temporal correlations between BOLD signals has been raised by multiple studies (see Murphy et al., 2013; Power et al, 2015 for reviews). In the current study, we conducted additional analyses using measures of head motion to ensure that movement was not different between L-dopa and placebo sessions, and that correlations between modulations in brain and behavior were not affected by overall head motion (see Figure S3). Furthermore, we ensured that the participants heart rate before, during, and after scanning were not significantly different between the first and second session or between L-dopa and placebo session (see Table S2).

#### Behavioral data

For each medication session, we calculated average response speed on correct trials (i.e., an inverse of response time) and a bias-free measure of perceptual sensitivity, d’ (MacMillan and Creelman, 2004). In order to assess the overall effect of L-dopa on behavioral performance across participants, we performed a separate repeated-measures analysis of variance (ANOVA) on response speed and perceptual sensitivity d’. To control for potential confounding factors in the interpretation of L-dopa effects, covariates of no interest (as listed above) were regressed out from the behavioral measures prior to conducting an ANOVA.

To visualize the robustness of L-dopa-induced change in individual participant’s performance (Figure 1B), single-subject 95% confidence intervals (CIs) were constructed based on bootstrapped performance differences between L-dopa and placebo. For each participant, we created 1,000 replications of the data by resampling the participant’s single-trial data of each session (i.e., resampling of 128 trials); based on these data we estimated the corresponding behavioral measure for each session. For a given behavioral measure, a significant change under L-dopa (versus placebo) per participant was inferred if the bootstrapped 95% CI did not cover zero.

#### Medication effects on brain measures

On the whole-brain level, statistical comparisons of signal variability and network metrics between L-dopa and placebo sessions were based on exact permutation tests for paired samples. We used Cohen’s d for paired samples as the corresponding effect size (Gibbons et al., 1993; Hentschke and Stuttgen, 2011; Lakens, 2013).

On the regional level, and for the analysis of both BOLD signal variability and network metrics, a bootstrap procedure with 10,000 replications was employed to test the medication effect. The bootstrap procedure allowed us to use a CI-based correction method to account for multiple comparisons across cortical regions (see below). For every cortical region per participant, we first computed the difference between the regional metrics (i.e., L-dopa–placebo). Then, the bootstrap distribution of the mean difference across all participants was estimated and used for statistical inference. Finally, Cohen’s d for paired samples was calculated as an effect size (Gibbons et al., 1993; Hentschke and Stuttgen, 2011; Lakens, 2013).

#### Correlational analyses

We further performed correlation analyses to examine whether the individual magnitude of L-dopa-induced modulations of BOLD signal variability and networks would relate to the individual magnitude of behavioral modulations. To this end, we computed the L-dopa-induced modulations by taking a difference (i.e., L-dopa–placebo) in both brain and behavioral measures. As such, we treated the placebo session as baseline for both brain and behavior during the auditory working memory task. Consistent with the analysis of behavioral data described above, we accounted for the potential confounding factors prior to the correlation analyses by regressing out the covariates of no interest (i.e., medication-session order, BMI, time from medication administration to scan onset).

Correlations between L-dopa-induced modulations in brain dynamics (i.e., BOLD signal variability or brain network metrics) and modulations in task performance were tested using rank-based Spearman correlation (rho). On the whole-brain level, the significance of the correlations was tested using a permutation test with 10,000 randomizations (Pesarin and Salmaso, 2010). On the regional level, a bootstrap procedure with 10,000 replications of the correlation coefficients was used for statistical inference. Using the same bootstrap procedures, we further examined the relationship between dopamine-related modulation of BOLD signal variability and modulations of the functional connectome.

#### Significance thresholds

For all statistical tests, we used α = 0.05 (two-sided) as uncorrected threshold of significance. For the regional analysis, and to correct for multiple comparisons entailed by the number of brain regions, we implemented the false coverage-statement rate (FCR) correction method (Benjamini and Yekutieli, 2005; applied e.g. in Obleser et al., 2010; Alavash et al., 2017). This method first selects the regions where the bootstrap distribution of the mean difference or correlation coefficients do not cover zero at the confidence level of 95%. In a second correction pass, FCR-corrected CIs for these selected regions are (re-)constructed at a level of 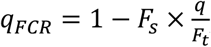, where *F_s_* is the number of selected regions at the first pass, and *F_t_* is the total number of brain regions. The tolerated rate for false coverage statements is *q*, here 0.05, at which the enlarged FCR-corrected intervals thus control the false-positive rate at the desired level.

## Results

### Effect of L-dopa on behavioral performances

Figure 1B illustrates the effect of medication (i.e., L-dopa vs. placebo) on overall behavioral performances in the auditory working memory task. Neither behavioral performance measures showed a consistent medication effect overall across participants (repeated-measures ANOVA with nuisance variables controlled; main effect of medication on response speed: F_1,21_ = 0.22, p = 0.65, η^2^_p_ = 0.01; main effect on perceptual sensitivity d’: F_1,21_ = 0.13, p = 0.73, η^2^_p_ = 0.006).

However, there was sizable inter-individual variation in the degree to which behavior benefitted (or suffered) from the L-dopa manipulation (see Cools and D’Esposito, 2011 for review). For both behavioral performance measures, we found about an equal split of participants showing opposite directions of the L-dopa effect (Figure 1B). It is possible that the direction of enhancement versus detriment from DA change can depend upon task performance levels (Cools and D’Esposito, 2011; Garrett et al., 2015; Gibbs and D’Esposito, 2005). To test whether the high individual variability is driven by the session at which L-dopa was administered, we examined a potential interaction effect of medication × medication–session-order. We did not find an effect for sensitivity d’ (F_1,17_ = 2.99, p = 0.10, η_p_^2^ = 0.15), but found a significant interaction effect on response speed (F_1,17_ = 6.15, p = 0.024, η_p_^2^ = 0.27), indicating that individuals receiving L-dopa on the second session exhibit greater behavioral improvement (t_20_ = 2.83; p=0.01) but to differing degrees. Hence, in the next sections, we further investigated whether such inter-individual differences of benefit from DA (see also Sharot et al., 2012) can be explained by L-dopa-induced modulations in brain dynamics. Since the focus of the study is to examine the effect of L-dopa alone, we examined the inter-individual differences while controlling the effect of session order.

### Modulation of signal variability under L-dopa vs. placebo

We started by investigating whether L-dopa modulates BOLD signal variability. We tested BOLD signal variability changes (L-dopa vs. placebo) on the whole-brain (i.e., average signal variability across all regions) as well as regional level. On the whole-brain level, a slight increase in BOLD signal variability with L-dopa relative to placebo was observed, but this increase was not statistically significant (M_L-dopa–Placebo_ = 1.33; Cohen’s d = 0.24; p = 0.30).

Next, we continued by analyzing signal variability on the regional level. L-dopa-induced modulations of signal variability were evident and statistically significant across a distributed set of sensory as well as heteromodal cortical regions. As illustrated in Figure 2A, we observed increased BOLD signal variability under L-dopa in bilateral superior temporal cortices extending from anterior to posterior portions, in bilateral inferior frontal gyri (IFG), and in right supramarginal gyrus (SMG). Also, L-dopa increased signal variability in visual cortical regions including bilateral fusiform gyri. In addition, an L-dopa-induced BOLD signal variability increase was observed in the cingulate cortex, bilateral motor cortices in the pre- and post-central regions, and the right temporo-parietal junction. Except for one region in the left middle frontal gyrus exhibiting a decrease in variability under L-dopa (Cohen’s d = 0.31), our results demonstrate that a distributed set of cortical regions relevant for the task increases its signal variability under L-dopa. On the contrary, applying the same contrast (i.e., L-dopa versus placebo) to mean BOLD activation during task performance (see Figure S7 for details) did not yield any region showing a significant medication effect.

**Figure 2.**
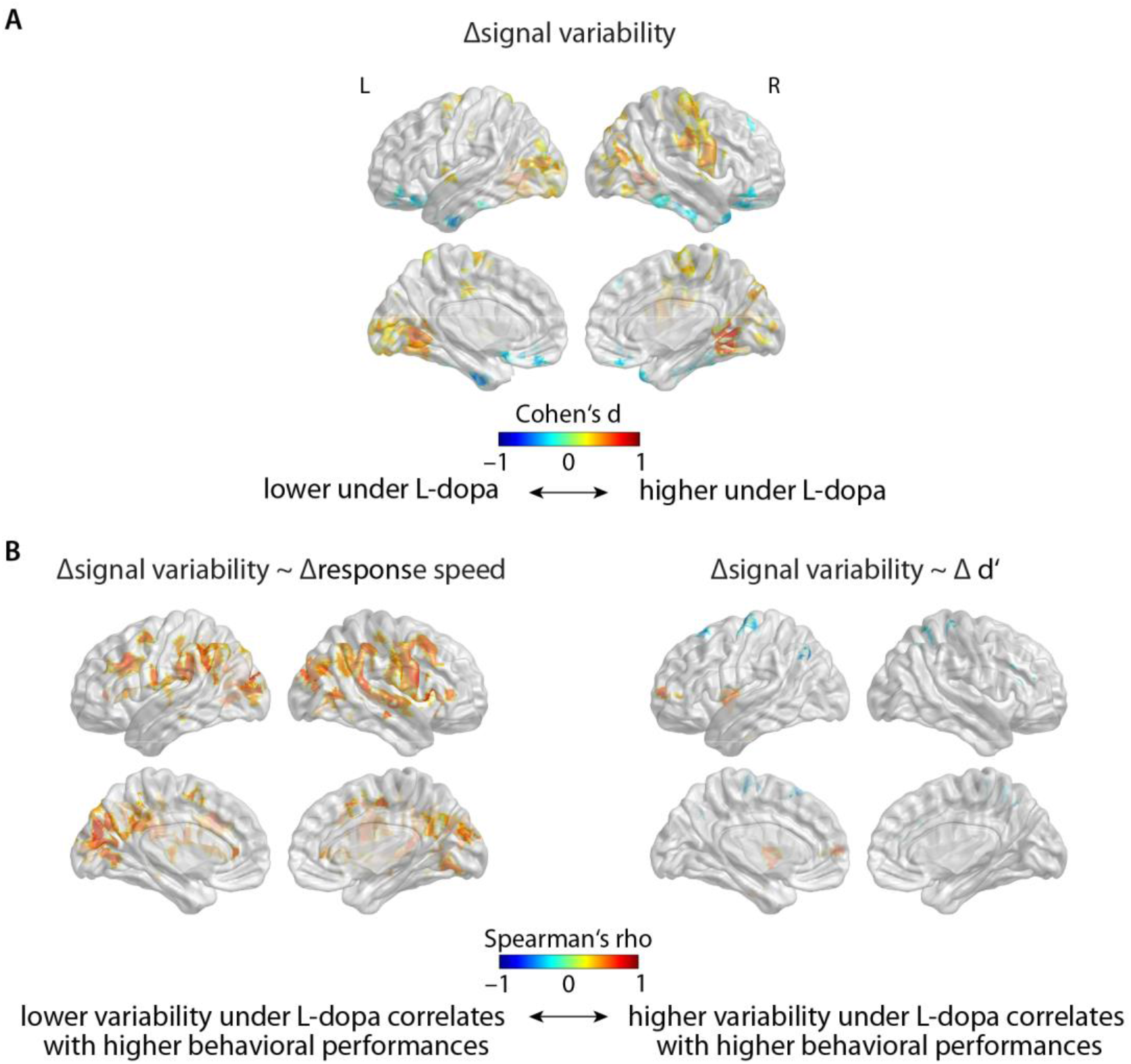
L-dopa-related modulation of BOLD signal variability and its relation to modulations of behavioral performance. **(A)** Cortical regions exhibiting significant change in signal variability under L-dopa versus placebo. The difference in signal variability (i.e., ∆: L-dopa–placebo) was expressed as Cohen’s d effect size. **(B)** Cortical regions exhibiting significant correlations between L-dopa-induced modulations of signal variability and behavioral performance. Response speed (left) and perceptual sensitivity d’ (right). Statistical significance (corrected for multiple comparisons) was ensured based on non-zero coverage of FCR-corrected 95% CIs of the 10,000 replications of bootstrapped distribution of the respective measures in (A) and (B) for each region (see Materials and Methods). Visualizations on the cortical surface were rendered using BrainNetViewer (Xia et al., 2013). L: left; R: right.

Subsequently, we investigated whether the extent of L-dopa-driven modulations in BOLD signal variability correlated with individuals’ behavioral benefits from L-dopa. For each region, we evaluated the relationship based on 10,000 bootstrapped iterations of correlation (i.e., Spearman’s rho) between modulations of BOLD signal variability and modulations of behavioral performance under L-dopa (i.e., L-dopa–placebo).

With respect to response speed, individuals’ BOLD signal variability modulations under L-dopa correlated with their gain in response speed under L-dopa. Specifically, greater signal variability in right IFG, bilateral precentral regions, bilateral angular gyri/SMG, and left parietal regions, correlated with faster responses under L-dopa (Figure 2B, left). The opposite pattern (i.e., lower variability correlating with higher response speed gain under L-dopa) was only observed in the subgenual anterior cingulate cortex.

With respect to perceptual sensitivity, the degree of BOLD signal variability modulation also correlated positively with L-dopa-induced change in d’—especially so in the left insula, bilateral superior temporal pole, and left parahippocampus (Figure 2B, right). Conversely, signal variability modulations in left postcentral/motor cortical regions exhibited significant negative correlations with d’ change. We further investigated the relationship between the elements of the change scores by examining the brain-behavior correlations separately under each placebo and L-dopa session, which did reveal strong modulations in one specific session (Figure S9).

### Whole-brain modulation of the functional connectome under L-dopa vs. placebo

One of the aims of the present study is to address how dopaminergic modulation impacts the brain network dynamics engaged in a cognitive task. We approached this question by measuring functional connectivity and integration of large-scale cortical networks using graph-theoretical metrics. The fMRI data were obtained from the participants who performed the auditory working memory task separately under L-dopa and placebo. We thus compared the brain network metrics between L-dopa and placebo across participants. This comparison revealed significant modulations in the connectivity and topology of brain networks under L-dopa versus placebo (Figure 3). The degree and direction of these modulations, however, were not homogenous across participants.

**Figure 3.**
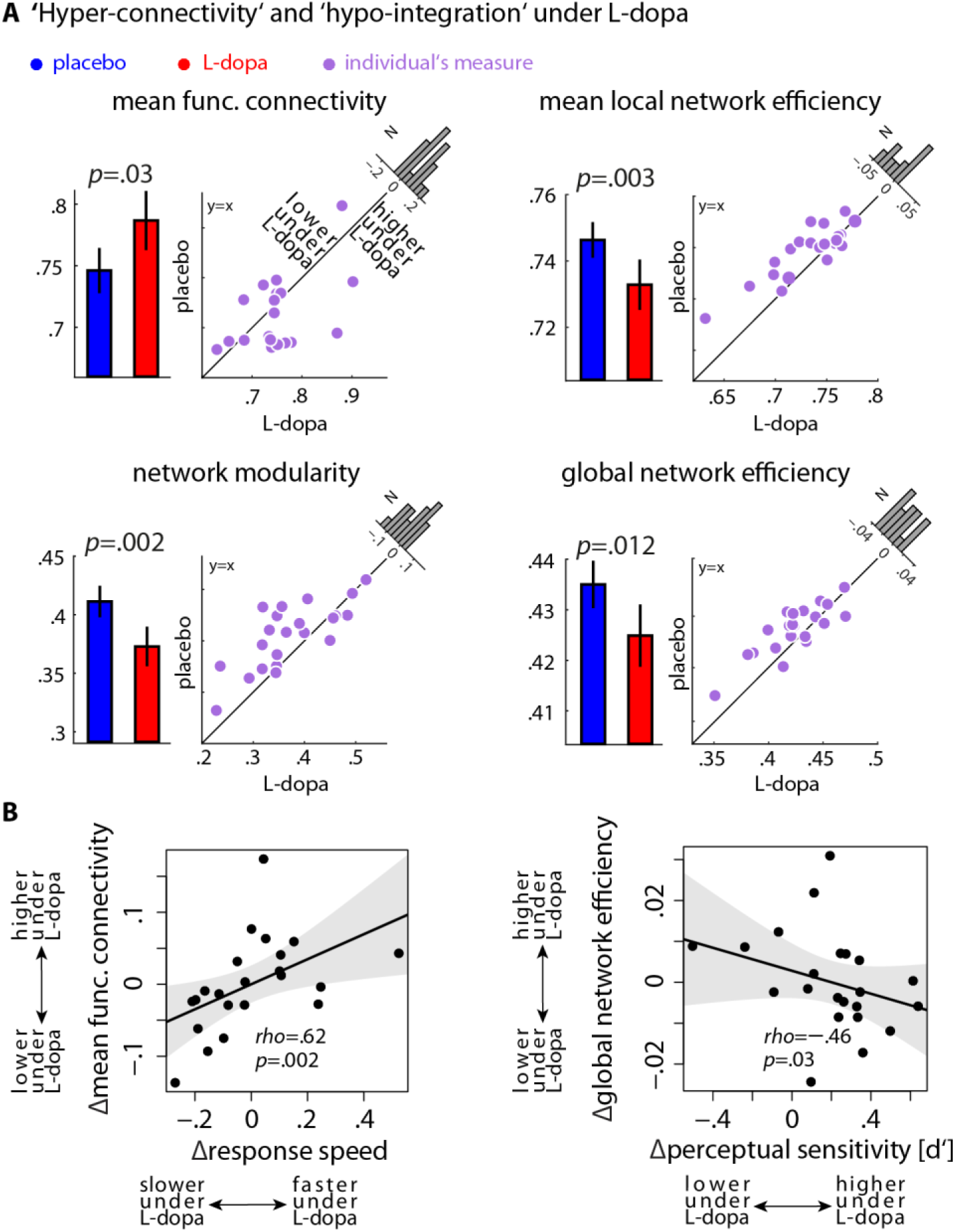
Modulation of the functional connectome under L-dopa and its relation to modulations of behavioral performance. **(A)** Whole-brain network metrics under L-dopa and placebo were compared using permutation tests. Bar plots show group average brain network metrics (error bars: ±1 SEM). Scatter plots illustrate the degree and direction of modulations (L-dopa versus placebo) for each participant. Histograms show the distribution of the modulations (L-dopa minus placebo) across all participants. Under L-dopa relative to placebo, 13/22 participants showed increase in functional connectivity—a relative ‘hyper-connectivity’. Moreover, in most of the participants functional integration of brain networks decreased under L-dopa—a relative ‘hypo-integration’—on all three topological scales: local network efficiency (14/22 participants), network modularity (15/22 participants) and global network efficiency (16/22 participants). **(B)** Correlations between modulations of the functional connectome and modulations in behavioral performance (∆: L-dopa–placebo). Participants who showed higher degrees of hyper-connectivity under L-dopa responded faster. Besides, participants with more pronounced hypo-integration under L-dopa (i.e., lower global network efficiency) showed higher perceptual sensitivity d’. The significance of the Spearman’s correlations (rho) was tested using permutation tests with 10,000 randomizations. Shaded area shows two-sided parametric 95% CI. Note that, prior to correlation analysis, all measures were residualized w.r.t. medication session order (L-dopa first versus placebo first), BMI, and time from medication administration to scan onset.

First, on the group level, functional connectivity showed a significant increase—that is, a relative ‘hyper-connectivity’—under L-dopa as compared to placebo (Cohen’s d = 0.4, p = 0.03; Figure 3A, top left). Across participants, the degree and direction of this modulation exhibited a sizable inter-individual variability: 13 out of 22 participants showed increase in functional connectivity. This relative hyper-connectivity was also observed when we compared the mean of raw functional connectivity strengths (i.e., without graph-thresholding; Cohen’s d = 0.35, p = 0.04). In addition, we found a significant alteration in the functional connectivity distribution—in particular its median and variance—under L-dopa relative to placebo (Figure S4).

Second, functional integration of brain networks on the group level showed a significant decrease under L-dopa versus placebo. This relative ‘hypo-integration’ of brain networks was consistently observed on the local, intermediate and global scales of topology, with a sizable inter-individual variability in the degree and direction of the modulation. More specifically, local efficiency of brain networks—a measure inversely related to the topological distance between network nodes within local cliques or clusters—was lower in 14/22 participants under L-dopa than placebo (Cohen’s d = 0.43, p = 0.003; Figure 3A, top right). In addition, modularity of brain networks—grouping of partner nodes within sub-networks—significantly decreased under L-dopa in contrast to placebo (Cohen’s d = 0.53, p = 0.002; Figure 3A, bottom left). Across individuals, network modularity was lower in 15/22 participants. Moreover, global network efficiency—a metric known to measure the capacity of brain networks for parallel processing—was significantly decreased under L-dopa compared to placebo (Cohen’s d = 0.4, p = 0.012; Figure 3A, bottom right). This relative hypo-integration was observed in 16/22 participants, and turned out to be—at least partly—due to higher fractions of disintegrated nodes in brain networks under L-dopa compared to placebo (Figure S1). Thus, L-dopa diminished network integration within the functional connectome on different scales of topology, with inter-individually differing degrees. Similar results were found when brain networks were constructed on different wavelet scales (Figure S5).

Knowing the heterogeneity of L-dopa-induced modulations in our sample, we next investigated whether the reorganization of the functional connectome under L-dopa explains the degree of the participants’ behavioral gain from L-dopa during the working memory task. We tested the correlation between the modulation (i.e., L-dopa–placebo) in the functional connectome on the one hand, and the modulation in task performance on the other hand. The change in task performance correlated significantly with both, the modulation in functional connectivity and global efficiency of brain networks (Figure 3B). Specifically, participants who showed higher degrees of hyper-connectivity under L-dopa also displayed higher gains in response speed under L-dopa than placebo (Figure 3B, left). Besides, participants with lower degrees of global network efficiency (equivalently more pronounced hypo-integration) under L-dopa showed higher perceptual sensitivity (d’; Figure 3B, right). However, the correlation between modulations in mean local efficiency or network modularity (*Q*) of the functional connectome and either of the performance measures were not significant (*E_local_* ~ response speed: rho = 0.01, p = 0.9; *E_local_* ~ d’: rho = –0.23, p = 0.3; *Q* ~ response speed: rho = − 0.26, p = 0.23; *Q* ~d’: rho = 0.19, p = 0.4). To examine the correlations between the elements that make up our change scores, we also investigated the brain-behavior correlations separately under each placebo and L-dopa session (Figure S9).

### Regional modulation of the functional connectome under L-dopa vs. placebo

To localize cortical regions contributing to the L-dopa-induced modulations observed on the whole-brain level (Figure 3), we analyzed the functional connectome on the regional level. To this end, we used nodal network metrics representing a given network property at each cortical region (see Materials and Methods: *Connectome analysis*). Consistent with the whole-brain level analysis, we compared the regional network properties of the functional connectome between L-dopa versus placebo. We found distributed cortical regions showing significant modulations in their functional connectivity and integration under L-dopa compared to placebo (Figure 4A).

**Figure 4.**
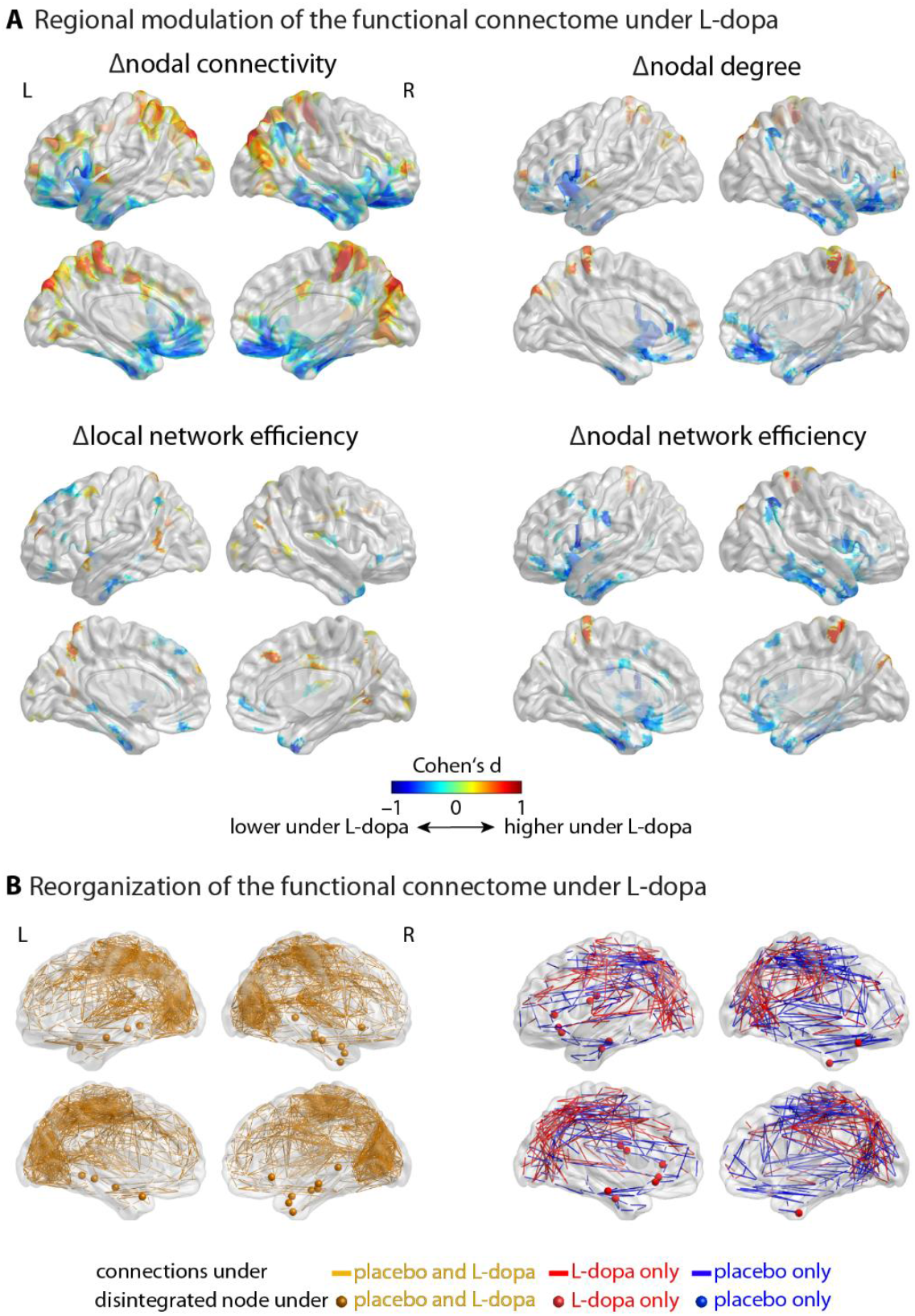
Cortical regions exhibiting functional connectome modulations under L-dopa. **(A)** Nodal network metrics of cortical regions were compared between L-dopa and placebo using bootstrap procedures with 10,000 replications of group average differences (∆: L-dopa–placebo). The significance of the differences was inferred based on the FCR-correction method accounting for multiple comparisons (p <0.05). Functional connectivity and network integration were significantly increased (warm regions) or decreased (cold regions) under L-dopa in contrast to placebo, with predominant hypo-integration of temporal and cingulo-opercular regions (cold regions). **(B)** Visualization of the topological reorganization of the functional connectome under L-dopa versus placebo. Orange links in the left panel represent putative functional connections engaged during the auditory working memory task under both placebo and L-dopa. A number of disintegrated nodes were consistently present under both placebo and L-dopa, mainly within the left and right temporal cortices (orange nodes). Blue links in the right panel represent functional connections which exist under placebo only. Red links represent functional connections which exist under L-dopa only. Compared to the core network (left panel), the functional connectome displayed an additional set of disintegrated nodes under L-dopa in temporal and cingulo-opercular regions (red nodes) but not under placebo (no blue node). Functionally disintegrated nodes were identified as nodes having a degree of zero. Group-average networks were constructed by thresholding the mean functional connectivity maps (network density set to 5% for better visualization). See Figure S2 for a thorough investigation of graph thresholding.

These modulations were found in both directions, i.e. L-dopa > placebo and L-dopa < placebo. More precisely, under L-dopa relative to placebo, regions overlapping mainly with paracentral lobule and precuneus underwent a relative hyper-connectivity (Figure 4A first panel, warm regions; Figure 4B, red connections). The most consistent pattern pointed to both hypo-connectivity and hypo-integration under L-dopa within temporal and cingulo-opercular regions (Figure 4A, cold regions; Figure4B, red nodes). In particular, nodal connectivity and nodal degree of cortical regions mostly in the inferior division of temporal cortex as well as frontal operculum were decreased under L-dopa relative to placebo (Figure 4A, first row). In addition, nodal network efficiency of regions within the inferior portion of temporal cortex, anterior cingulate and frontal operculum was significantly lower under L-dopa than placebo. These findings were further supported by the existence of higher number of disintegrated nodes under L-dopa versus placebo (Figure 4B, red nodes). Similar results were found when a functional parcellation was used to construct the brain graphs (Figure S8).

Next, we tested the correlation between regional modulation (i.e., L-dopa–placebo) of the functional connectome and modulations in behavioral performances. We found significant correlations between modulations of behavioral performances and modulations of nodal connectivity and nodal efficiency (Figure 5). Consistent with the correlations observed on the whole-brain level (Figure 3B), participants who showed increased nodal connectivity (i.e., hyper-connectivity) in bilateral superior temporal/angular gyri and anterior cingulate under L-dopa, also responded faster under L-dopa (Figure 5, left, red circles). Moreover, decreased nodal network efficiency (i.e., hypo-integration) of a brain node within the right frontal operculum/insula under L-dopa correlated with higher behavioral gains in perceptual sensitivity (Figure 5, right, blue circle).

**Figure 5.**
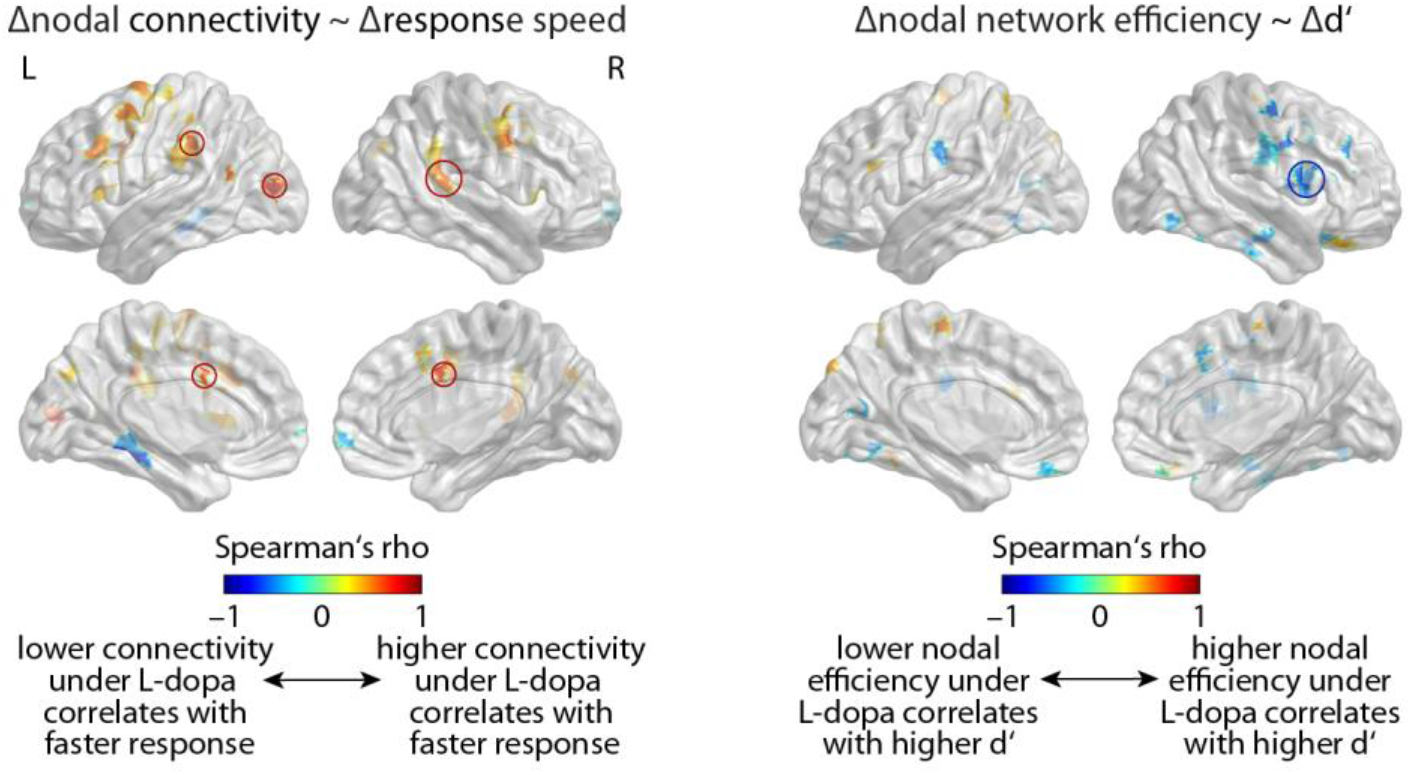
Cortical regions where modulation of the functional connectome correlated with modulations in task performance. Modulation of nodal connectivity in bilateral superior temporal/angular gyri and anterior cingulate (left panel, red circles) under L-dopa positively correlated with the change in response speed during the auditory working memory task (∆: L-dopa–placebo). Additionally, lower nodal network efficiency of a brain node within the right frontal operculum/insula (right panel, blue circle) under L-dopa correlated with higher perceptual sensitivity d’. The correlation at each region was tested using a bootstrap procedure for Spearman’s rho (10,000 replications). Circles highlight the regions that survived FCR correction (p < 0.05). Colored regions without circles are visualized at the uncorrected level.

### Relationship of modulations in signal variability and modulations in the functional connectome

Given that L-dopa significantly modulated BOLD signal variability and network organization of the functional connectome, we further investigated whether these modulations in cortical dynamics directly relate to each other across participants. Specifically, we examined which cortical regions exhibit a relationship between the modulation of their signal variability under L-dopa (versus placebo) and the modulation in their network configuration within the functional connectome under L-dopa (versus placebo). We thus separately analyzed the correlations between the modulations of three regional network metrics (i.e., nodal connectivity, local network efficiency, and nodal network efficiency) and modulation in signal variability across cortical regions.

Figure 6 illustrates cortical regions exhibiting robust correlations between modulations in BOLD signal variability and regional network metrics under L-dopa (versus placebo). Across participants, we found that the extent of L-dopa-induced changes in BOLD signal variability positively correlated with the extent of nodal connectivity changes in broadly distributed cortical regions, spanning temporal, cingulate, frontal, and parietal cortices (Figure 6, top).

**Figure 6.**
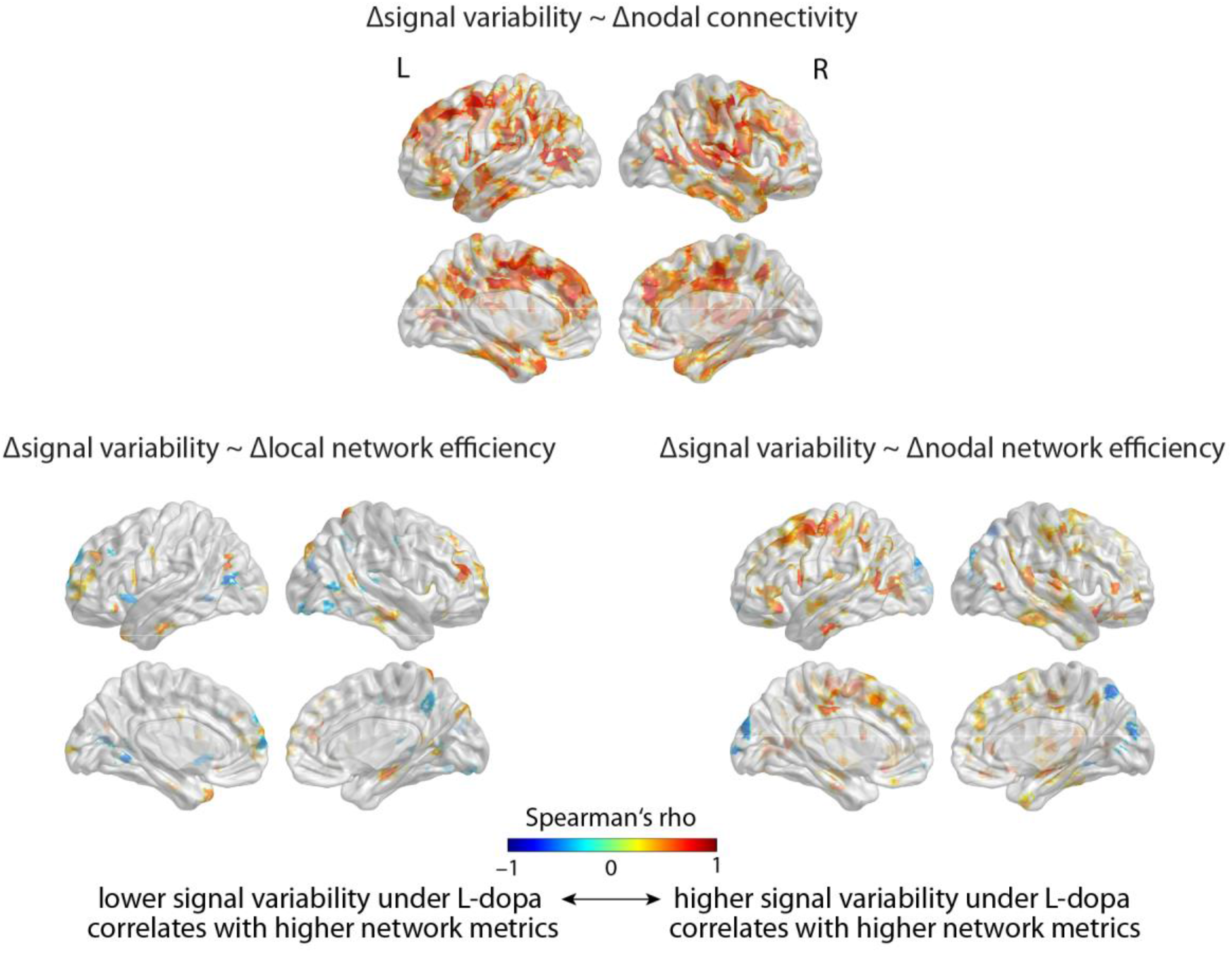
Cortical regions exhibiting significant correlations between the respective modulations of BOLD signal variability and the functional connectome under L-dopa. The extent of L-dopa-induced modulation (∆: L-dopa– placebo) of signal variability showed predominantly positive correlations to that of nodal connectivity (top), local network efficiency (bottom, left) and nodal network efficiency (bottom, right) across participants. The significance of the correlations were inferred based on the FCR correction accounting for multiple comparisons (p < 0.05).

Interestingly, L-dopa-related modulation of BOLD signal variability correlated with the extent of modulation in individuals’ brain network efficiency, both on local and global scales of topology. Specifically, network efficiency within each node’s neighborhood graph (i.e., local efficiency) as well as its functional integration to the whole-brain graph (i.e., nodal efficiency) significantly correlated with BOLD signal variability in distributed cortical regions. Increased signal variability specifically in temporal, left inferior frontal, cingulate, as well as insula cortices correlated with higher network efficiency in similar regions (Figure 6, bottom). Notable exceptions were visual areas (within the occipital and inferior temporal cortices), which showed negative correlations between modulations in BOLD signal variability and network efficiency.

### Modulation of striato-cortical connectivity

Knowing that the midbrain dopaminergic system innervates striatal and cortical networks (Jaber et al., 1996; Seger and Miller, 2010; Frank, 2011), we further investigated two follow-up questions. First, we asked which striatal structure would show a significant modulation in its functional connectivity with the cortical network under L-dopa relative to placebo. Second, and more specifically, we asked *which* cortical nodes would show modulation in their functional connectivity with the striatal structure(s). To address these questions, we used a functional parcellation of the striatum with seven seed regions (Choi et al., 2012) from which we extracted their mean hemodynamic signals (wavelet scale 2: .06–.12 Hz), and computed their respective correlation strengths with all cortical nodes. Finally, to capture the strongest correlations, the top 25% strongest connections were selected and averaged per striatal seed (mean nodal connectivity; similar to Choi et al. (2012)). Prior to the correlation analysis, and in order to account for physical proximity of the striatum to the insula and orbital frontal cortices, we regressed out the mean signal of the cortical voxels that were up to 9 mm away from the left and right putamen (similar to Choi et al. (2012)).

We found that, under L-dopa relative to placebo, a striatal seed region in the posterior ventral putamen showed higher nodal connectivity with the cortical network (p<0.05; Figure 7A). This region included ten 2×2×2-mm striatal voxels, which were mainly within the right striatum (MNI slice coordinates: coronal y=–18, axial z=–6). A similar result was found when the cortical nodes were defined based on the functional parcellation (Cohen’s d=0.67; p<0.05, FCR-corrected) or when different connectivity thresholds were used in the range 5%–30% in steps of 5%. However, the correlation between this modulation and the modulation in behavior was not significant (d’: rho=0.16, p=0.47; response speed: rho=0.26, p=0.24).

**Figure 7.**
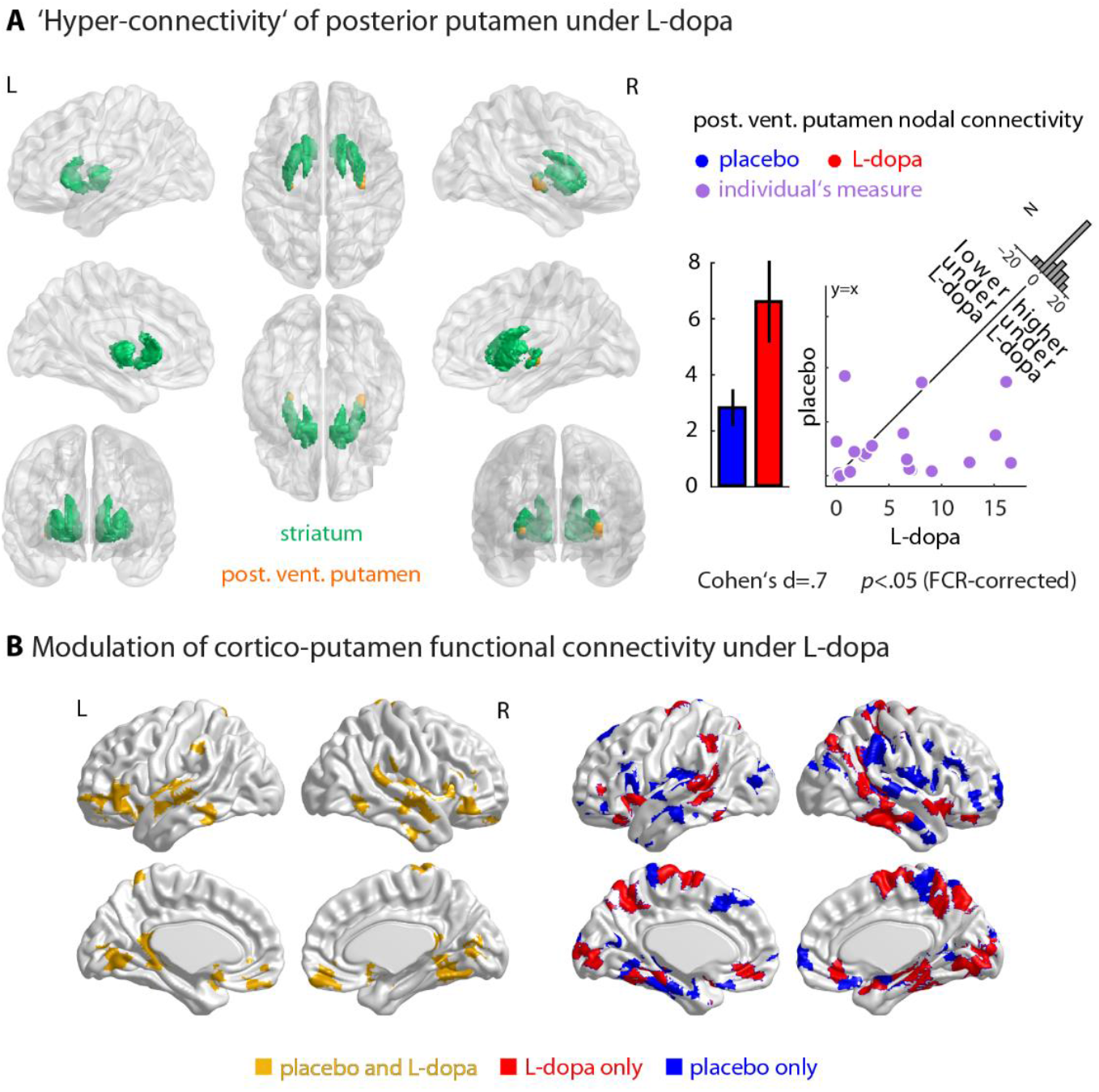
Modulation of striato-cortical functional connectivity under L-dopa relative to placebo. **(A)** Hemodynamic signals were extracted from seven striatal seed regions using a functional parcellation, and their one-to-all cortical correlation strengths were compared between L-dopa and placebo using bootstrap procedures. A striatal seed region at the posterior ventral putamen (shown in orange) displayed higher nodal connectivity with the cortical network under L-dopa in 17/22 participants (MNI slice coordinates: coronal y=– 18, axial z=–6). Scatter plots illustrate the degree and direction of modulations (L-dopa versus placebo) for each participant. Histograms show the distribution of the modulations (L-dopa minus placebo) across all participants. **(B)** Group-averaged functional connectivity map between each cortical node and the posterior putamen seed region found in (A) was constructed by keeping the top 25% strongest correlations. The cortical regions which were functionally connected with the posterior putamen are color-coded and visualized separately for placebo, L-dopa or both.

Subsequently, to map the cortical consequence of the posterior putamen hyper-connectivity under L-dopa, we constructed the group-averaged functional connectivity map between each cortical node and the posterior putamen seed region by keeping the top 25% strongest correlations. This procedure was done separately for each placebo and L-dopa session. The resulting cortical map is shown in Figure 7B. We observed that cortical regions mostly overlapping with the supplementary motor area, ventromedial prefrontal, as well as intraparietal lobule and bilateral temporal cortices showed more extended functional connectivity with the posterior putamen under L-dopa relative to placebo (Figure 7B, red). With respect to striatal signal variability, we only found a significant decrease of variability in the caudate head under L-dopa relative to placebo (FCR-corrected p<0.05; Cohen’s d=0.46). This region is shown in red in Choi et al. (2012), and has been associated with the default mode network on the cortical level, including the posterior cingulate and medial prefrontal cortices.

## Discussion

How does dopamine modulate large-scale cortical dynamics, and how does this in turn relate to differences in cognitive performance among individuals? We addressed this question by investigating the effect of L-dopa on BOLD signal variability and the functional connectome while young healthy adults performed an auditory working memory task. We examined L-dopa-induced modulations of brain dynamics and behavior relative to placebo, and the relationships between these modulations across participants.

### Heterogeneity of L-dopa effects across individuals

In our sample, we did not observe a main effect of L-dopa on behavioral performance, but a notably high inter-individual variability in benefits from L-dopa. Previous accounts suggest that the absence of an overall effect may be due to the well-known inverted U-shaped effect of DA: Individuals’ baseline levels of cognitive performance, endogenous DA, and/or genetic variations in dopamine metabolism determine the direction of L-dopa-related benefits/detriments (Goldman-Rakic et al., 2000; Vijayraghavan et al., 2007; Cools and D’Esposito, 2011; Pearson-Fuhrhop et al., 2013). Another source of variability is that weight-dependent dosage may explain the degree of benefits from L-dopa found in old participants (Chowdhury et al., 2012). Interestingly, however, the present study with a sample of young healthy participants does not show any significant linear or quadratic trends related to the weight-dependent L-dopa dosage on task performance. In parallel to the previous accounts, our results provide new insights into the role of dopamine in the context of working memory, and suggest that heterogeneity of L-dopa-induced effects might also arise from differences in individuals’ brain signal variability and functional network architecture, as it is discussed in the following sections.

### L-dopa increases BOLD signal variability

In our sample, L-dopa increased BOLD signal variability on the group level. Notably, the individual extent and direction of this modulation correlated with the behavioral benefit from L-dopa. Our findings lie in parallel with recent studies emphasizing the importance of signal variability in predicting well-functioning cognitive operations (Garrett et al., 2010; Grady and Garrett, 2014), and the role of DA in boosting BOLD signal variability (Garrett et al., 2015), and further illustrates that DA effects across individuals vary according to their baseline hemodynamic signal variability.

Specifically, we found that L-dopa increased signal variability in distributed cortical regions deemed relevant for the auditory working memory task (see Figure S7). These regions include bilateral superior temporal cortices, often associated with auditory and speech perception (Liebenthal et al., 2005; Obleser and Eisner, 2009). A similar effect was also observed in the bilateral IFG, implicated in pitch and verbal processing and working memory (e.g., Griffiths, 2001; Gaab et al., 2003; Zatorre et al., 1994). A separate analysis of mean BOLD activation did not reveal any region showing a significant difference between mean BOLD activations under L-dopa versus placebo on the corrected level (Figure S7). These results indicate that BOLD signal variability reflects a different aspect of neural computation as compared to event-related mean BOLD activation.

Interestingly, distinct cortical regions appeared relevant in explaining individuals’ behavioral benefits from L-dopa. Our results show that loci of signal variability modulations that correlated with response speed versus d’ gains are indeed different. Signal variability in motor and parietal regions seems to underlie response speed gain, whereas L-dopa-induced sensitivity benefits were associated with signal variability changes in parahippocampal area, where activity is related to episodic memory/encoding (Amaral and Insausti, 1990; Davachi et al., 2003; Chowdhury et al., 2012). As we have not observed speed–accuracy trade-offs with the benefits from L-dopa (p = 0.90), it is a tenable hypothesis that signal variability in distinct cortical regions contributes complementarily to behavioral performance.

The current study is limited in that we only measured these BOLD signal properties in response to L-dopa. But it is important to note that the changes in signal variability due to L-dopa may reflect underlying complex changes in neurovascular coupling and brain metabolism: while L-dopa administration enhances signal-to-noise ratio of neural activity and increases cerebral blood flow (CBF), it decreases BOLD response (Zaldivar et al., 2014).

### L-dopa substantially modulates the functional connectome

During the auditory working memory task, L-dopa reduced the functional integration across cortical regions on the group level. This hypo-integration manifested at different topological scales, and overlapped with temporal and cingulo-opercular regions. Since the brain graphs were matched in network density across participants and sessions, this hypo-integration cannot be due to network density differences (cf. van den Heuvel et al., 2017). Further, the L-dopa-induced hyper-connectivity and hypo-integration were consistent across a wide range of network densities at different graph thresholds (Figure S2). Disintegration of certain nodes, mostly in temporal and cingulo-opercular regions, may be interpreted as diminished statistical dependency (co-activation or functional cross-talk; not to be mistaken by anatomical disconnection) between these and other cortical regions during auditory working memory processing. Thus, this reorganization most likely stems from a functional disintegration of temporal and cingulo-opercular nodes from regions less relevant for the task (arguably the default mode network; see Guitart-Masip et al., 2016; Nagano-Saito et al., 2017).

Note that we use the terms ‘integration’ and ‘disintegration’ within the framework of graph theory. In a functional connectome and on the global scale of network topology, for a brain graph to display greater network integration (increased capacity for parallel processing), long distance correlations are required. In sparsely connected functional networks (as in our case), short distance connections predominate, and are typically associated with greater strength of connectivity (Vértes et al., 2012). Thus, increase in connectivity most likely would indicate formation of local neural processing (equivalently, lower functional integration, Rubinov and Sporns, 2010). As such, a network shifts from having a globally-integrated topology to a more clustered and segregated topology. In our data, however, we observed a significant decrease in global efficiency, but also in local efficiency and modularity (see Figure S10 for the correlations across network metrics). Accordingly, the network reorganization was not a functional shift toward higher segregation. We thus investigated an alternative functional reorganization, namely nodal disintegration, which could potentially attenuate all three topological measures. Our analysis revealed significantly stronger nodal disintegration under L-dopa compared to placebo (Figure S1).

Interestingly, the above hypo-integration correlated with L-dopa-induced sensitivity benefits: participants with more pronounced hypo-integration on both whole-brain and regional levels (right frontal operculum/insula) showed higher d’. These correlations were specific to network efficiency— a measure inversely related to topological distances, which further supports the functional disintegration of task-relevant regions.

Accordingly, L-dopa-induced hypo-integration might indicate a functional shift of the temporal and cingulo-opercular regions from a globally integrated state to a more autonomous engagement in task processing. Previous work suggests that the cingulo-opercular network is crucial for sustained attention (Dosenbach et al., 2006, 2007; Vaden et al., 2013), hemostatic salience processing (Seeley et al., 2007), tonic alertness (Sadaghiani and D’Esposito, 2015; Coste and Kleinschmidt, 2016) and dynamic coordination between the default mode network and the central executive network (Menon and Uddin, 2010; Uddin, 2015). Thus, the more autonomous state of this network can increase the signal-to-noise ratio in cortical information processing, and help maintain to-be-attended (here, syllable pitch) information. The other potentially relevant interpretation of L-dopa-induced network disintegration is that L-dopa preferentially supports the phasic dopaminergic receptor (D1) network state responsible for stable maintenance of representations with high signal-to-noise ratio while suppressing the flexible network state (i.e., D2-state) that allows processing of distractor or task-irrelevant neural activity (see Durstewitz and Seamans (2008) for computational models). However, this possibility needs further testing as it is yet unclear whether and how overall increase of presynaptic dopamine levels (i.e., via L-dopa) differentially activates D1- and D2-class receptors.

To further investigate whether the effects arise from task-related neural dynamics or modulation of spontaneous activity (Ponce-Alvarez et al., 2015; Bolt et al., 2017), we used a general linear model (GLM) with finite impulse response (FIR) basis functions (0–22 s relative to trial onset) to estimate task-related BOLD activity. Subsequently, the residual data obtained from the GLM was used for analyzing signal variability and connectivity of spontaneous activity (Zhang and Li, 2010; Cole et al., 2014; Alavash et al., 2015b; Finc et al., 2017; Bolt et al., 2017). This analysis did not reveal any significant difference in signal variability or network dynamics of spontaneous brain activity between placebo and L-dopa. This result supports that, in our sample, L-dopa modulated task-related cortical dynamics, thereby altering the signal variability and the functional connectome of cortical regions engaged during the auditory working memory task.

To date, only few studies have investigated the effect of DA on brain networks, mostly focusing on resting-state or fronto-striatal functional connectivity (Nagano-Saito et al., 2008; Kelly et al., 2009; Cole et al., 2013; Bell et al., 2015). Our study, in contrast, focuses on large-scale cortical interactions during a listening task with attention and memory components. On the whole-brain level, we found an overall hyper-connectivity under L-dopa. On the regional level, however, we found that L-dopa decreased nodal connectivity of temporal and cingulo-opercular regions (Figure 4, cold regions and red nodes), whereas it increased nodal connectivity of paracentral lobule and precuneus (Figure 4, red connections). This apparent inconsistency between the direction of the global and regional results can be explained by network disintegration. We found that temporal and cingulo-opercular regions were functionally disintegrated (Figure 4B; red nodes) from the remaining of the whole-brain network (Figure 4B; red connections). The connected component includes the paracentral/precuneus nodes, whose significant increase in nodal connectivity contribute to the hyper-connectivity observed on the whole-brain level. The disintegrated nodes, on the other hand, do not contribute to whole-brain functional connectivity simply due to the absence of connections (hypo-connectivity/-integration). Thus, functional connectivity on the whole-brain level remained significantly higher under L-dopa relative to placebo, the extent of which correlated with individuals’ response speed gains from L-dopa.

### Modulation of signal variability correlates with modulations of the functional connectome

DA-driven modulation of signal variability showed significant positive correlations with modulations of nodal connectivity and network efficiency mainly in temporal and anterior cingulate cortices known to be crucial for listening behavior (Langers and Melcher, 2011; Erb et al., 2013). One might associate these findings with the fact that connectivity between two signals, expressed as their correlation, requires a certain amount of variability. However, the present findings more likely reflect a systematic relationship between two measures of cortical dynamics. First, our findings point to the correlation between modulations (L-dopa versus placebo) in two measures of brain dynamics rather than within-session correlations. Second, regional network metrics are multivariate measures: nodal connectivity is estimated as mean correlation between a region’s signal and its node-pairs’ signals, and nodal network efficiency is estimated based on topological distances between a given node and every other node in the network. Notably, for two regional signals to show coherent fluctuations over time, not only variability (information) but also covariance (coordination) is necessary (Carbonell et al., 2014; Cole et al., 2016). Thus, correlation between the modulation in variability of a regional signal and the modulation in the network wiring of the same region is most likely a systematic relation between two measures of cortical dynamics.

Furthermore, such systematic relationship in our data was supported by the direct linear relationship between functional connectivity and variability which was only found to be positive (Figure 6A). Taken together, our study motivates the importance of investigating both aspects of cortical dynamics—that is, signal variability and connectivity—as two complementary measures.

These findings can be understood in terms of information-processing capacity (McIntosh et al., 2008; McIntosh et al., 2014), and integration of information across the functional connectome (Tononi et al., 1994; Tononi et al., 1998; Sporns, 2003; Stam and van Straaten, 2012). Our findings are consistent with previous studies reporting positive correlations between entropy of resting-state EEG signals and nodal centrality and network efficiency (Mišić et al., 2010, 2011). Nodes conveying more information are characterized by many connections and short/direct paths to the rest of the network, thereby optimally ‘relaying’ local information across the connectome (see Timme et al., 2016 for a recent evidence based on networks of neural spikes). More recently, Garrett et al. (2017) showed that signal variability of thalamic activity up-regulates the functional integration of brain networks as measured by network dimensionality. Overall, our findings suggest that L-dopa modulates cortical signal variability as well as dynamic communication engaged in processing task-relevant information, and the extent of these modulations can be a mechanistic determinant of the often-observed inter-individual differences in behavioral benefits from L-dopa.

### L-dopa modulates posterior putamen cortical connectivity

Previous studies have emphasized the importance of striatal dopamine in working memory (Frank et al., 2001; Cools et al., 2008; Landau et al., 2009; Guitart-Masip et al., 2016), and even observed an opposite effect of signal variability change with aging in cortical versus subcortical structures (e.g., Garrett et al., 2010). In our sample, we found that posterior putamen cortical connectivity was significantly altered under L-dopa, with 17/22 participants showing a hyper-connectivity relative to placebo. According to Choi et al. (2012), the posterior ventral putamen represents the dorsal attention network when functionally mapped onto the cortex. These results thus provide insights into how striatum might contribute to our findings, and suggest that L-dopa-driven hyper-connectivity of the functional connectome is likely to arise, at least partly if not entirely, from higher posterior putamen cortical modulation.

### Conclusion

We showed that an increase of dopamine level while performing an auditory working memory task modulated BOLD signal variability and the functional connectome predominantly in temporal and cingulo-opercular regions. Notably, the degree of these modulations predicted inter-individual differences in behavioral benefit from L-dopa. The findings thus provide insights on understanding the source of prominent inter-individual differences of L-dopa effects on cognitive performance. Our study fills a critical gap in knowledge regarding dopaminergic modulation of univariate and multivariate aspects of cortical dynamics. The data provide first evidence for a direct link between dopaminergic modulation of BOLD signal variability and the functional connectome. Dopamine thus appears to maintain the dynamic range of, and communication between, cortical systems involved in processing task-relevant information during cognitive performance.

## Acknowledgement

Research was supported by the Max Planck Society (Max Planck research grant to JO) and the European Research Council (ERC Consolidator grant AUDADAPT, no. 646696, to JO). We thank Sarah Tune (University of Lübeck) for her assistance in statistical analysis and for stimulating discussions.

